# Kinetic network model to explain gain-of-function mutations in ERK2 enzymes

**DOI:** 10.1101/409482

**Authors:** M. Misiura, A. B. Kolomeisky

## Abstract

ERK2 is a kinase protein that belongs to a Ras/Raf/MEK/ERK signalling pathway, which is activated in response to a range of extracellular signals. Malfunctioning of this cascade leads to variety of serious diseases, including cancers. This is often caused by mutations in proteins belonging to the cascade, frequently leading to abnormally high activity of the cascade even in the absence of external signal. One such *gain-of-function* mutation in ERK2 protein, called a *sevenmaker* mutation (D319N), was discovered in 1994 in Drosophila. This mutation leads to disruption of interactions of other proteins with D-site of ERK2 and results, contrary to expectations, in increase of its activity *in vivo.* However, no molecular mechanism to explain this effect has been presented so far. The difficulty is that this mutation should equally negatively affect interactions of ERK2 with *all* substrates, activators and deactivators. In this paper, we present a quantitative kinetic network model that gives a possible explanation of the increased activity of mutant ERK2 species. A simplified biochemical network for ERK2, viewed as a system of coupled Michaelis-Menten processes, is presented. Its dynamic properties are calculated explicitly using the method of first-passage processes. The effect of mutation is associated with changes in the strength of interaction energy between the enzyme and the substrates. It is found that the dependence of kinetic properties of the protein on the interaction energy is non-monotonic, suggesting that some mutations might lead to more efficient catalytic properties, despite weakening inter-molecular interactions. Our theoretical predictions agree with experimental observations for the *sevenmaker* mutation in ERK2. It is also argued that the effect of mutations might depend on the concentrations of substrates.

## INTRODUCTION

Mitogen-activated protein (MAP) kinase ERK2 (Extracellular Signal-Regulated Kinase 2) is an enzyme that plays important role in a variety of biochemical processes. It is activated in response to several extracellular signals such as mitogen, interleukin, growth factors and cytokines (1,2), operating as a part of a Ras/Raf/MEK/ERK signalling pathway, which is crucial for cell functioning (3-5). ERK2 a small 42 kDa protein, consisting of C-terminal and N-terminal domains (6-8). It’s an ATP-dependent enzyme and the ATP-binding site, as well as the catalytic site, are located in the region between the main domains. Unlike many enzymes, ERK2 does not bind its substrates in the immediate vicinity of catalytic site, but instead it utilizes the so-called recruiting sites (docking sites), which are located 15-20 Å away from the place where the catalysis occurs. These binding sites are usually referred to as D-recruiting site (DRS) and F-recruiting site (FRS), and they are responsible for recognition of multiple substrates with different structures (9-19).

To become catalytically active, ERK2 requires phosphorylation of two of its residues: Tyrl85 and Thrl83 (20). Phosphorylation leads to alteration of mutual orientation of domains and their dynamics (7, 21-25%). Activation of ERK2 is normally done by MAP/ERK kinases (MEK) (26). Active ERK2, in its turn, can be deactivated by a number of phosphatases (1, 20). Combination of these two processes - activation and deactivation - enables precise control of ERK2 activity, providing a robust and efficient method to respond to external signals. Since ERK2 regulates many critically important processes, including cell growth, cell differentiation and cell proliferation, the alteration of its normal enzymatic activity can lead to serious negative effects, such as uncontrollable tissue growth, which was shown to be linked to a variety of diseases, including cancers (27-30).

An interesting example of ERK2 malfunctioning is the existence of gain-of-function mutations inside the Ras/Raf/MEK/ERK signalling pathway. Such mutations can alter the structure of one of the kinases in a phosphorelay, thus preventing the activity of ERK2 from being regulated properly, and eventually leading to disease (3, 30). Hie most known gain-of-function mutation in ERK2 (D319N) is called *sevenmaker,* and was it discovered in Drosophila in 1994 as a result of genetic screenings for mutations that activate the sev signalling pathway in the absence of signal (31-34). The mutation is located in the DRS (docking site) of ERK2 in the common domain (CD) region (35).

The fact that the *sevenmaker* mutation activates the enzyme is rather surprising, because it is expected that this mutation should negatively influence interactions of ERK2 with *all* substrates, activators and deactivators in a similar fashion. One would suggest then the mutation should lower the enzymatic activity. Indeed, there are experimental observations (35, 36) suggesting that many substrates, activators (including MEK) and deactivators, use the DRS site and, in particular, the CD domain to recognize the ERK2 protein. They interact using a so called kinase interaction motif (KIM), which consists of 2-3 positively charged Lys and/or Arg residues (19). Thus the *sevenmaker* mutation should disrupt all ERK2-involving processes in the similar way, so it is surprising that it can lead to apparent increase of ERK2 activity *in vivo* (31, 32). Despite the fundamental importance of ERK2, molecular mechanisms of its gain-of-function mutations (and specifically the *sevenmaker* mutation) remain not well understood. One proposal to explain these observations is based on the fact that there are only two ERK2 activators, MEK1 and MEK2, while there are many deactivators. It was suggested that some deactivators might be less, affected by the disrupted interaction with the CD domain (35). To support this, there are experimental data showing that the mutation D319N in the ERK2 is less sensitive to dephosphorylation (37, 38). However, that does not resolve the problem entirely since the ability of ERK2 to phosphorylate substrates should be also reduced by comparable amount (35).

In this paper, we propose a theoretical model that quantitatively explains the effect of the *sevenmaker* mutation. It is based on the kinetic network description of the system with the additional assumption that the mutation equally changes interaction energies between ERK2 and all substrates, activators and deactivators. By analyzing a simplified regulation network of ERK2, built as a system several coupled Michaelis-Menten processes, the kinetic properties of ERK2 proteins are evaluated explicitly via a first-passage method. It is shown that the effective chemical kinetic properties in these systems might change non-monotonically as a function of the interactions. This suggests that some mutations might lead to more efficient catalytic properties of ERK2 protein variants, despite the decrease of the interaction energies. It is argued that this is a possible molecular mechanism of gain-of-function mutations in ERK2, explaining the experimental observations on the *sevenmaker* mutation.

## METHODS

### Kinetic Network Model

To clarify the molecular mechanisms of increased activity for the *sevenmaker* mutation, one should analyze the kinetic properties of the biochemical regulation pathway of ERK2. Although it is known that ERK2 functioning involves many biochemical states and transitions, we consider a minimal simplified version of the regulation scheme as presented in Figure 1. Our idea is to approximate the regulation network as three coupled Michaelis-Menten processes that correspond to main processes involving this enzyme: activation, inactivation and ERK2-mediated phosphorylation (1). In the state 0 (labelled as ERK2) the enzyme molecule is inactive (not phosphorylated) and it can bind MEK enzyme with a rate constant *u* to reach the state 1 (labelled as ERK2-M): see Figure 1. From the state 1, ERK2 can return back to the state 0 with a rate *w* by dissociating from the complex with MEK, or it can be phosphorylated to reach the state 2 (ppERK2) with a rate *α.* After that, ppERK2 can either be dephosphorylated through the formation of a complex with a phosphatase (state 3, ERK2·D) with the rate constant *u,* or it can remain active and phosphorylate its own substrates by forming the substrate-enzyme complex with the rate constant *u* (state 4, ppERK2-S), and producing the product (state 5, ppERK2+P) with the rate *α.* To simplify calculations, in this model we assume that the corresponding rate constants in all Michaelis-Menten reactions for different processes are equal to each other so that there are only three kinetic parameters in the system: *u, w* and *α*. This assumption is based on the fact that all enzymatic processes are taking place at the same location and they involve chemical species that are not very dissimilar. But it is also important to notice that the relaxing of this condition (making all corresponding rates different) will not qualitatively change the main theoretical predictions of this work while it will make the mathematical calculations much more complicated.

**Figure 1:**
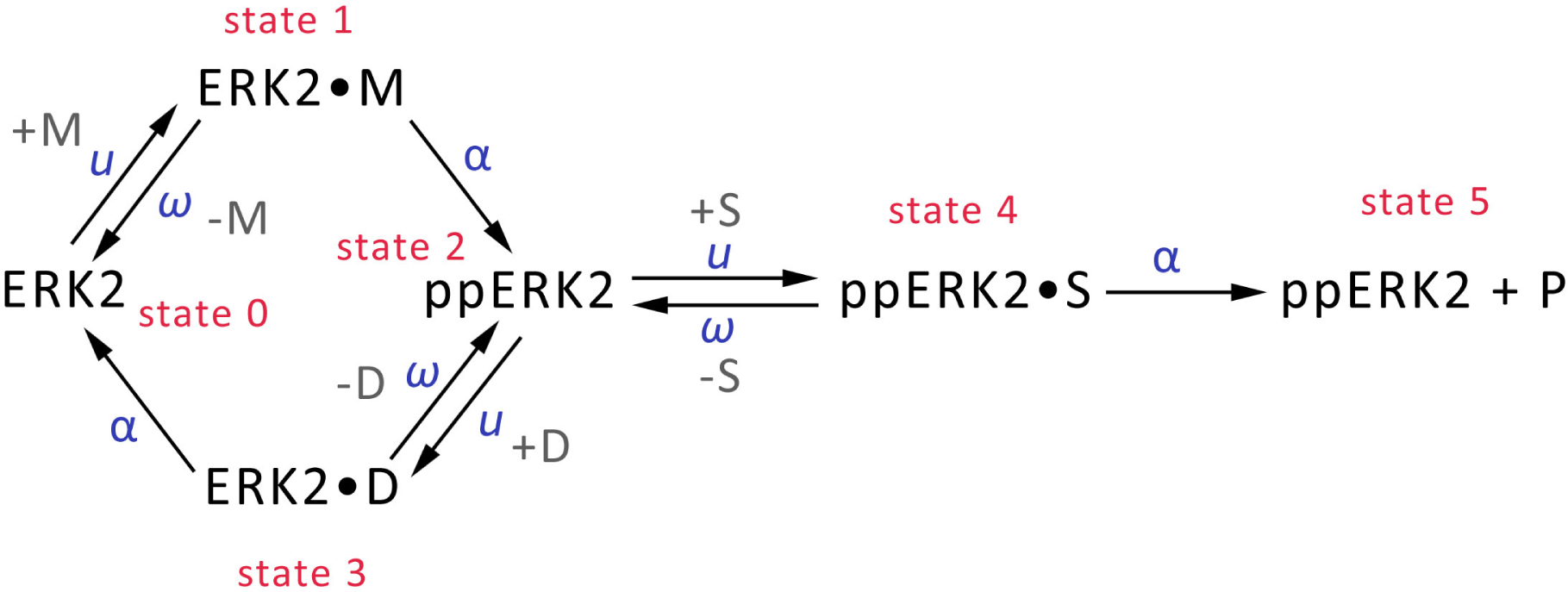
A simplified biochemical regulation scheme for ERK 2 considered in this work. ERK2 first must be phosphorylated by MEK (denoted as *M)* to become an active enzyme and to phoshorylate its substrates (denoted as *S*). At the same time, the phosphatase (denoted as *D*) can dephosphorylate ERK2 to return it to the inactive state. More details are in the text.

In evaluating the catalytic properties of this system, we employ a method of first-passage processes that was successfully utilized for analyzing multiple processes in Chemistry, Physics and Biology (39,40). The idea is to introduce a first-passage probability density function *F*_*n*_*(t),* which is defined as a probability to complete the reaction (i.e., to reach the final state 5) at time *t* if at *t* = 0 the system was in the state *n.* Determining these functions will provide a full dynamic description of the catalytic process in this system. The temporal evolution of first-passage probabilities is governed by a set of the backward master equations (39,40), which are closely related to standard chemical kinetics equations:

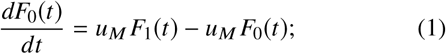

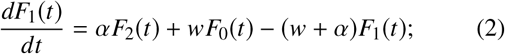

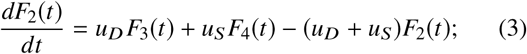

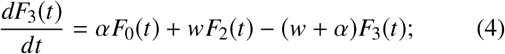

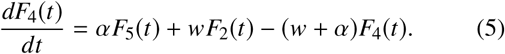

In these equations, we take into account the fact that the association transition rates are proportional to the concentrations of participants, i.e.

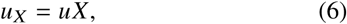

where *X = M, D* or *S* are the concentrations of activator, deactivator and substrate, respectively. In addition, the initial condition requires that *F*_5_(t) = *δ*(t), which means that if the system starts in the state 5 the reaction is accomplished immediately.

To calculate the first-passage probabilities, we utilize Laplace transformations, 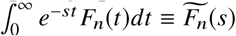. Then Eqs. (l)-(5) can be rewritten as simpler algebraic expressions:

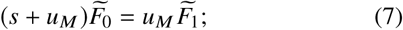

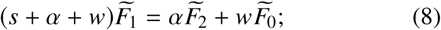

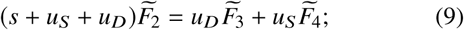

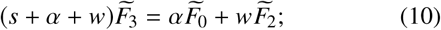

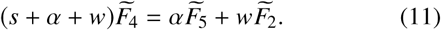

The initial condition also leads to 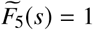. This system of equations can be easily solved. Specifically, for starting the process in the state 0 we obtain,

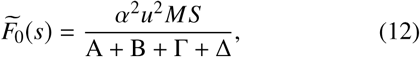

where new parameters are defined as

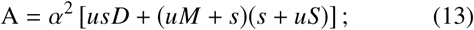

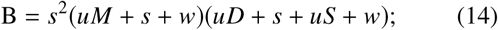

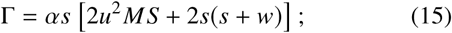

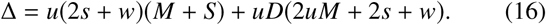

The explicit expressions for the first-passage probability functions provide a direct way of describing all dynamic properties in the system. For example, the average time to reach the product state 5 starting from the state 0, which is the same as the mean time for the catalytic reaction (turnover time) is given by (39, 40)

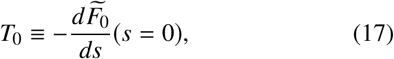

from which using Eqs. 12-16 we get

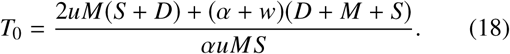

This result can be better understood if we rewrite it in the Michaelis-Menten-like form with respect to the substrate *S* transformation (*T*_0_=1/*k*_*cat*_ + *K*_M_*/k*_*cat*_ *1/*S*) as follows,

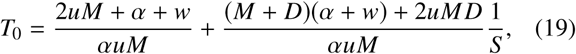

from which the overall effective Michaelis-Menten parameters for the kinetic network are determined in terms of the microscopic transition rates:

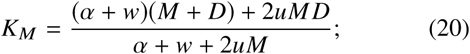

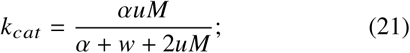

and

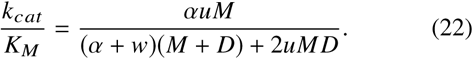

To quantitatively analyze the effect of mutations, we assume that mutations change the strength of the interactions in the ERK2 complexes with activators, deactivators or substrates, respectively. We define a binding energy ɛ as a measure of strength of such interaction. The sign is chosen so that more negative values of ɛ correspond to stronger binding. Then the detailed balance-like arguments allow us to estimate the relations between the rate constants and the binding energy:

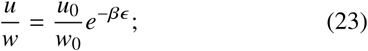

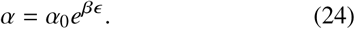

Here the rates with superscript 0 correspond to transition rates for the hypothetical situations when the interactions energies are equal to zero. These equations can be understood in the following way. The stronger the binding interactions, the faster the system will go into the states with the complex formation (states 1, 3 and 4), and the slower it will leave these states. Correspondingly, weaker interactions stimulate the system to preferentially break these complexes faster than to form them.

Determining the enzymatic properties of the system requires explicit expressions for rates that include the effect of the interactions. Then we can rewrite the expressions for the transition rates as (40):

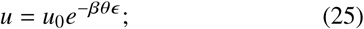

And

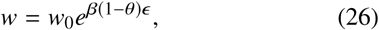

with *β= 1/k*_*B*_*T.* The parameter 0 ≤*θ* ≤ specifies how the interaction energy is distributed between forward and backward transitions to form or to break the complex state (40). For simplicity, in the following expressions we omit the subscript 0, so that *u, w* and *α* now replace *u*_0_, *w*_0_ and *α*_0_, respectively. With these assumptions, our final equations for the kinetic parameters are given by:

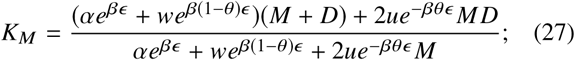

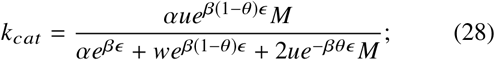

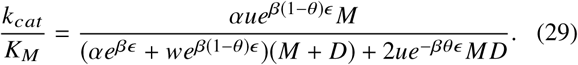

The main advantage of this theoretical approach is that now the effect of mutations can be investigated quantitatively because in our language it corresponds to varying the interaction energy *ɛ*.

### Analysis for Michaelis-Menten Model

To understand better the mechanisms of the ERK2 regulation that couple together several enzymatic processes, it should be compared with the simplest situation that involve only a single enzymatic process. For this purpose, we present here a brief derivation of catalytic properties for a classical Michaelis-Menten kinetic scheme:

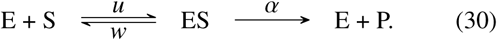

The derivation follows exactly the same steps as was described for the model in Figure 1 above, and only main steps are presented. We assume here that *E* + *S* corresponds to the state 0, *ES* describes the state 1, and *E* + *P* is the final state 2. Hie temporal evolution of the corresponding first-passage probability functions follows from

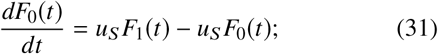

and

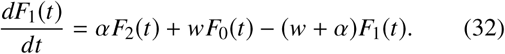

After the Laplace transformation, these equations can be rewritten as follows:

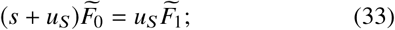

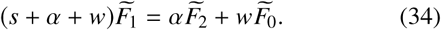

Solving this system of equations, yields the following expression for the turnover time *T*_0_:

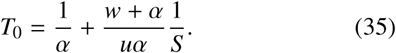

Finally, the Michaelis-Menten parameters are given by

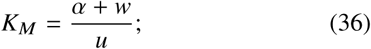

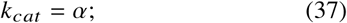

and

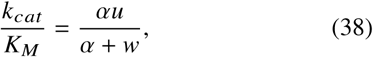

where *u. w* and *α* depend on the substrate binding energy exactly as described above [see Equations 24 - 26],

Unless stated otherwise, the following parameters are utilized for calculations in the Results and Discussion section: θ = 0.5, *α* = *w*· = 100*s*^-1^,*k =* 10,000*s*^-1^*M*^-1^, *D* = *M* = *S* = 0.001M. These parameters are chosen just to illustrate our theoretical method.

## RESULTS AND DISCUSSION

Our main idea is that the mutations modify the interaction energy between the enzyme and the substrate molecules, and this leads to changes in the chemical kinetic properties of the system. Using explicit expressions derived in the previous section, we can analyze how the enzymatic parameters for ERK2 and simple Michaelis-Menten (MM) schemes vary with the binding energy. The results are presented in Figures 2 and 3. One can see that the enzymatic properties of ERK2 regulation network differ significantly from the classical MM scheme. Lowering the strength of binding interactions (making ɛ more positive) strongly increases the catalytic rate *k*_*cat*_ in the MM system, while the dependence of *k*_*cat*_ on ɛ is non-monotonic for the ERK2 system (see Figure 2). It can be shown also that in tliis case the highest value of *k*_*cat*_ is achieved for

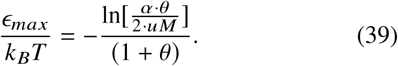

Varying the interaction energy also leads to different curves for the Michaelis constant for the simple MM and for the ERK2 regulation network (Figure 3). *K*_M_ strongly increases with ɛ in the MM case, while for the ERK2 system *K*_M_ is slowly changing between two limiting behaviors. For very strong attractive interactions *(ɛ*→- ∞), we have *K*_M_≃*D* while for strong repulsive interactions *(ɛ*→ +∞) *K*_M_≃*D* + *M.*

**Figure 2:**
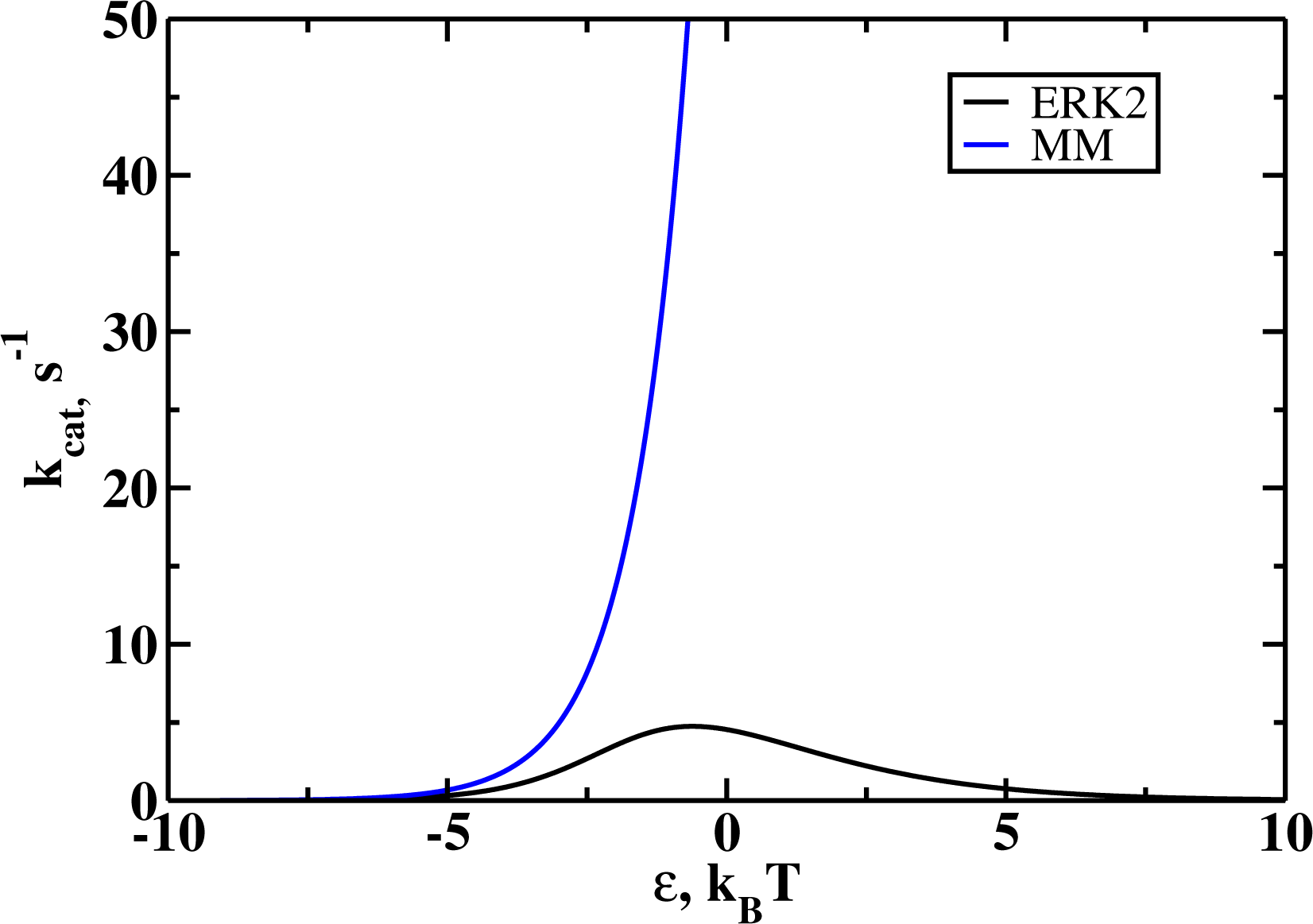
Catalytic constants as functions of the binding energies for Michaelis-Menten (MM, blue) and ERK2 schemes. Negative energies correspond to stronger binding. For MM scheme a monotonic behavior is observed: the stronger the interaction, the lower *k*_cat_ while for ERK2 scheme the dependence is non-monotonic: there is an optimal value of binding energy that produces the highest *k*_cat_.

**Figure 3:**
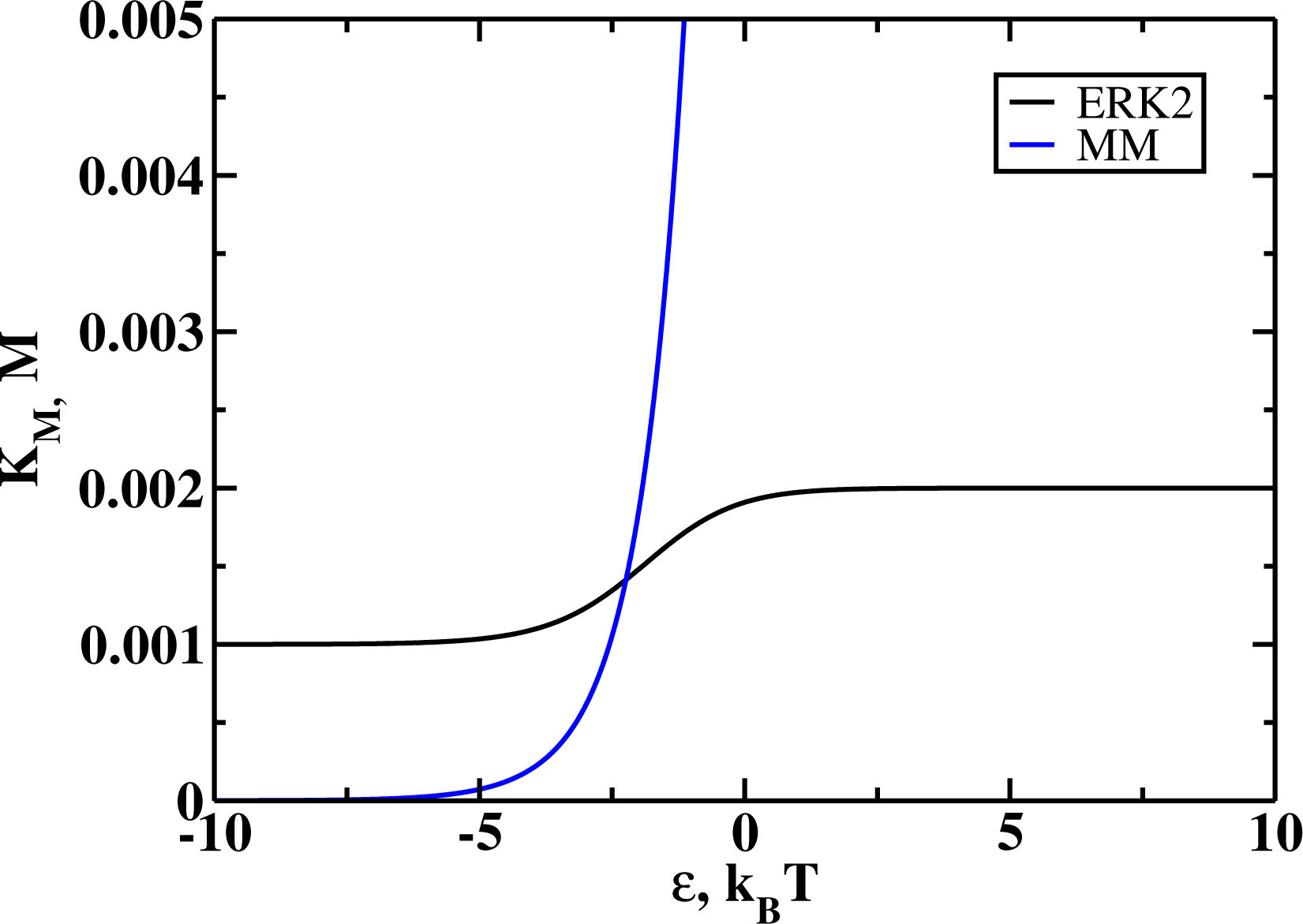
Michaelis constants as functions of the binding energies for Michaelis-Menten scheme (MM, blue) and ERK2 scheme (black). Negative energies correspond to stronger binding. For MM scheme the dependence is monotonic, and the stronger the binding, the lower the Michaelis constant. For ERK2 scheme the dependence is also monotonic, but *K*_m_ changes between two limits.

To quantify better the enzymatic efficiency of ERK2 proteins, it is more useful to consider a ratio *k*_*cat*_*/K*_M_, which is known as a specificity constant. The larger this parameter, the more efficient is enzymatic process. Figure 4 presents specificity constants as functions of the binding energies for both schemes, and again the classical MM behavior is strikingly different from the predictions for the ERK2 regulation system. The specificity constant for the MM process decreases monotonically with the interaction energy, while the non-monotonic dependence is observed for the ERK2 case. The position of the maximum here is

**Figure 4:**
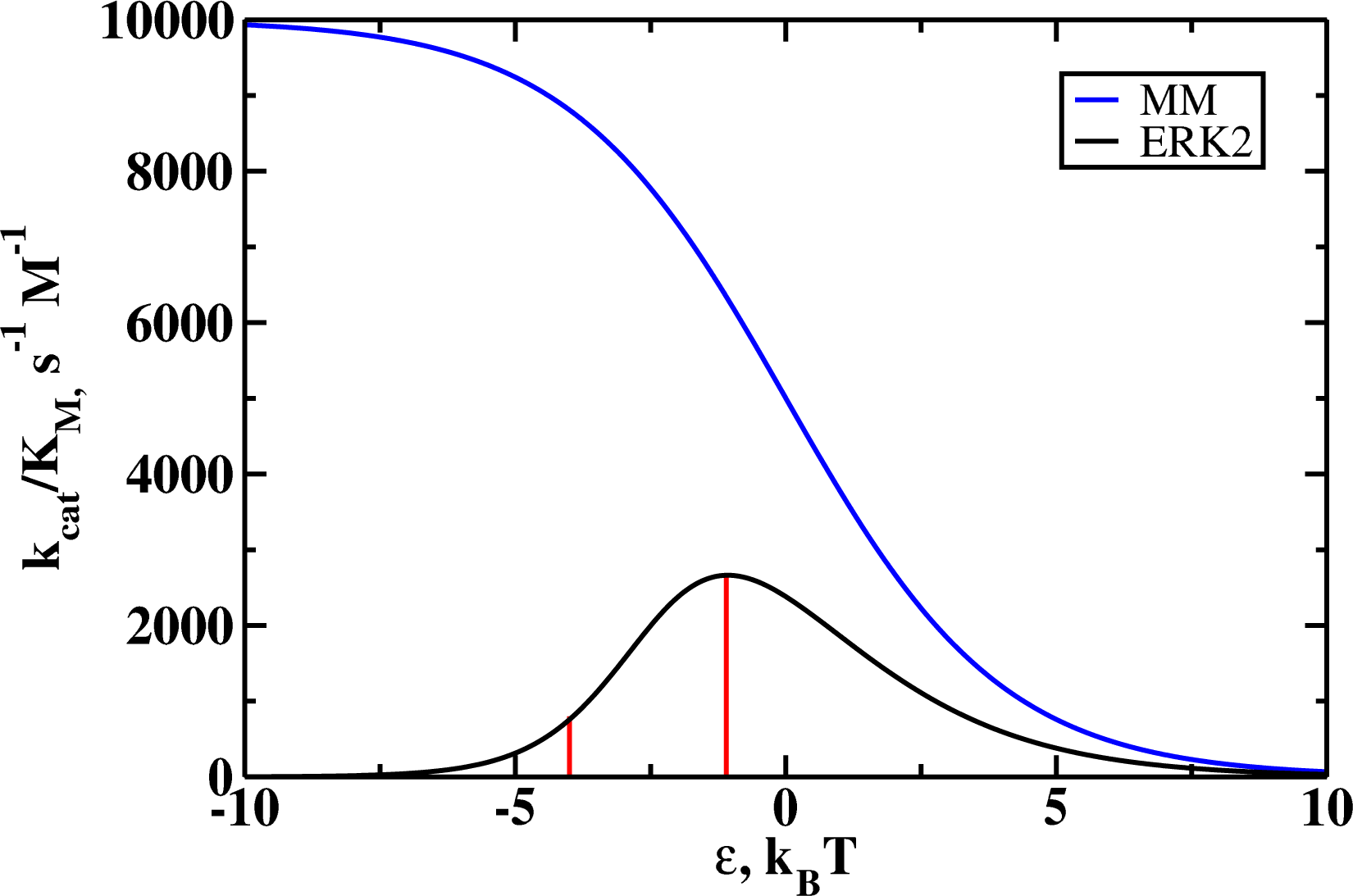
Specificity constants (*k*_*cat*_*/K*_m_) as a functions of the binding energy for the Michaelis-Menten scheme (MM, blue) and for the ERK2 scheme (ERK2, black). Negative energies correspond to stronger binding. In the region between two vertical lines for the ERK2 scheme the decrease of strength of interactions will lead to higher values of specificity.

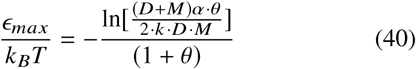

This result has a very important consequence for explaining the appearance of gain-of-function mutations in the ERK2 system. If one assumes that the binding energy in the WT enzyme (*ɛ* _WT_) is negative and it does not correspond to ɛ _max_ (*ɛ* _WT_ *< ɛ* _max_), then mutations that change the interaction energies to the range between ɛ *WT* and ɛ _max_ will increase the activity of enzyme: the region between two vertical lines on Figure 4. In this situation, the mutation that weakens the interactions with the substrate will effectively make the ERK2 regulation network more efficient in comparison with the wild type case. This might be a possible molecular mechanism of how the *sevenmaker* mutation operates in the ERK2 system. It is also important to note that since ERK2 is a regulatory enzyme it is likely to operate *in vivo* at low concentrations in the regime where the specificity constant is the main property that determines the catalytic efficacy.

The effect of gain-of-function mutations can be also explained using the fluxes along the different branches of the regulation scheme presented in Figure 1. The flux that starts in the state 0 reaches the state 2 via the activation branch (*J*_*a*_). where it divides into the flux that goes to the final product via the phosphorylation branch (*J*_*p*_,) and the flux that returns back to the state 0 via the deactivation branch (*J*_*d*_). In the stationary state, the flux balance requires that

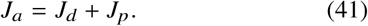

The overall enzymatic activity can be correlated with the product formation flux *J*_p_. Then our theoretical picture suggests that the *sevenmaker* mutation lowers both *J*_a_ and *J*_d_ fluxes, but it decreases the deactivation flux more so that the product formation flux *J*_p_ in the case of mutation is larger in comparison with the WT ERK2 molecule, i.e., *J*_p_*(mutant*) > *J*_P_*(WT).* The results presented in Figure 5, where the effect of varying the deactivation flux is investigated, support these arguments. Lowering the concentration of deactivator (D) decreases the possibility for the system to go into the deactivation branch. For low *D* the enzymatic properties of the ERK2 regulation pathway, as expected, approach the simple MM scheme, and the non-monotonic behavior as well as the ability to increase the enzyme’s activity by mutation disappear. Only when there are significant fluxes via the deactivation path the gain-of-function mutations might appear in such systems. Thus, the gain-of-function mutation in the ERK2 regulation network is the result of coupling of several enzymatic processes that work in opposite directions.

**Figure 5:**
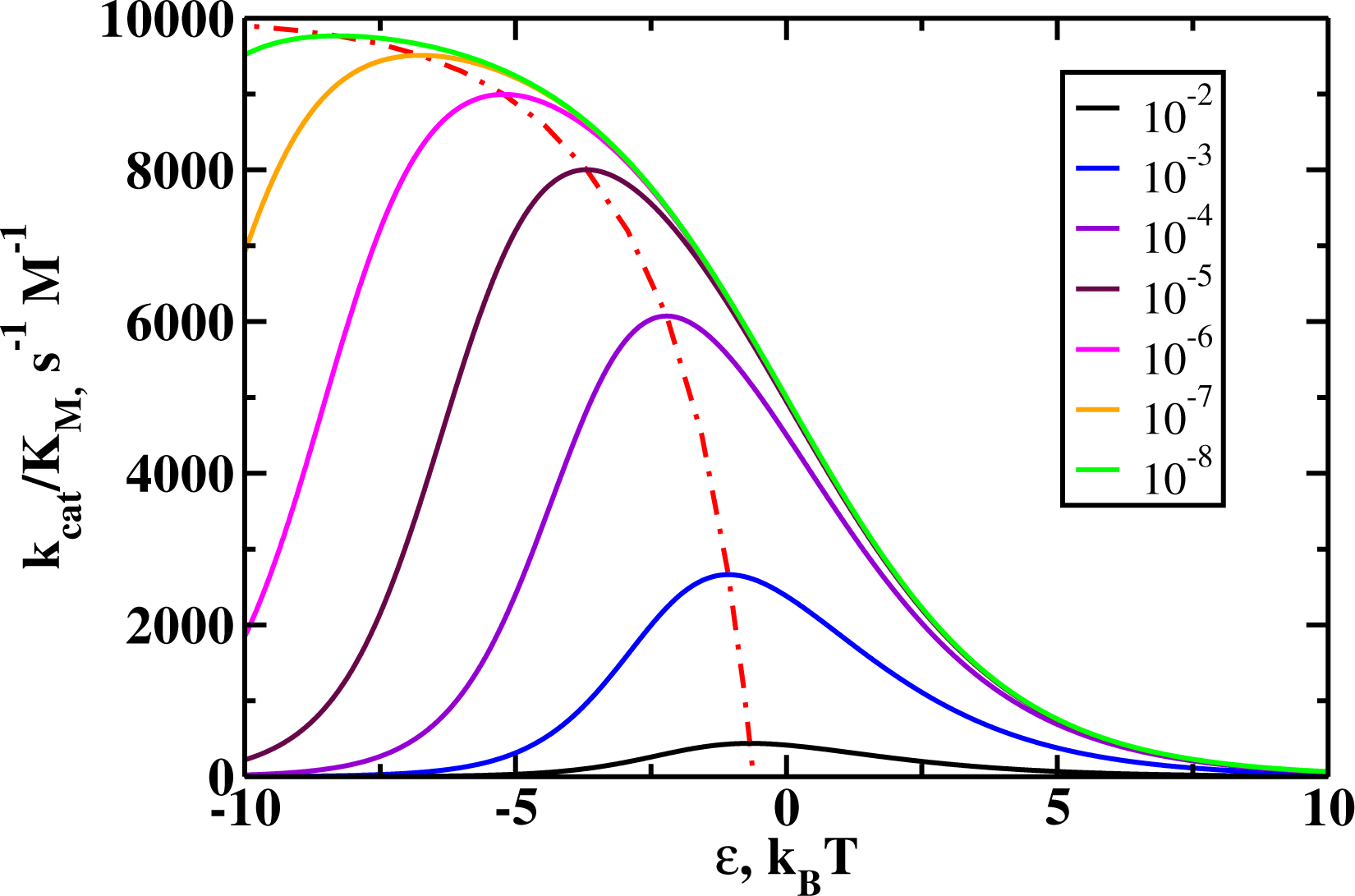
The specificity constant as the function of the interaction energy for varying contributions from the deactivation process. Numbers in the legend show the concentrations of the deactivator *D* in units of *moles/l.* Dashed line shows the position of the maximum of the specificity constant.

Theoretical calculations presented in Figure 5 also lead to a surprisinng prediction that the sign of the mutation effect (positive gain-of-function, or negative loss-of-function) can be reversed by changing the concentrations of the network components (activators, deactivators and substrates). If one assumes that the *sevenmaker* mutation operates in the range of interaction strengths as given in Figure 4 (between two vertical lines), then for very low concentrations of deactivators this mutation will no longer be increasing the enzymatic activity.

Because ERK2 has multiple substrates, we speculate that this ability of the network to selectively affect the efficiency of enzymatic processes might be an additional level of regulation that can benefit cellular functioning.

Our theoretical views can be further supported by analyzing the turnover times as a function of the interaction energies, as illustrated in the Figure 6. One can see that the effective overall catalytic rate (inverse turnover times) shows the non-monotonic behavior for both the simple MM and the ERK2 regulation pathways. But there is a range of interaction energies where the increase in the binding energy lowers the rate of the MM process, while the process in the ERK2 regulation network can go faster. This is an addition argument to explain the existence of the gain-of-function mutations and specifically effect of the *sevenmaker* mutation in ERK2. Even if the mutation lowers the rate along each enzymatic pathway, it might effectively increase the overall rate in the complex ERK2 scheme that combines all of them.

**Figure 6:**
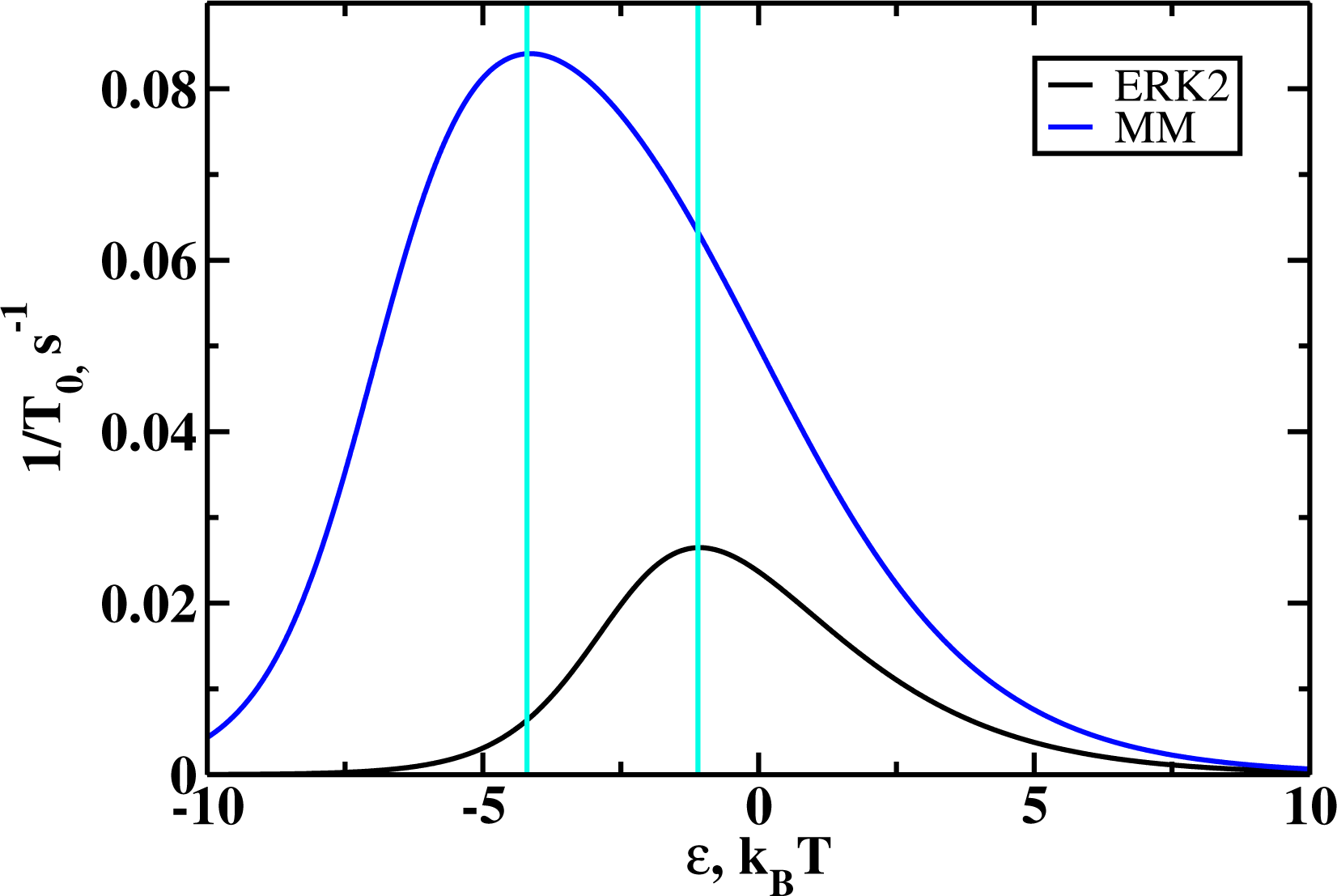
Inverse catalytic turnover times, or the effective overall reaction rates for the product formation, as a function of the interaction energy for the Michaelis-Menten scheme (MM, blue) and for the ERK2 scheme (black). Concentration of substrate is *S*=10^-5^. In the region between two vertical lines decrease of interaction energies leads to decrease of the reaction rate for MM scheme, but for the ERK2 scheme it leads to the increased turnover rate.

Because our theoretical approach makes quantitative predictions, it is important to compare them with experimental observations. However, experimental data on *sevenmaker* mutation are pretty scarce, mostly qualitative and obtained using very different techniques and under different experimental conditions. This prevents us from explicitly including them into our analysis. But we notice that Camps et al. (41) found that about one order of magnitude higher concentrations of MKP-3 are needed in order to deactivate the mutated *ERK2*^D319*N*^ protein variant as compared with the wild type *ERK2.* In addition, the decrease of deactivation activity by other phosphatases (PAC1, MKP-1 and MKP-2, approximately from 3 to 7 times lower) for mutated ERK2 species was reported by Y. Chu et al. (37). All these observations are consistent with our theoretical flux arguments. Furthermore, experimental data by T. Tanoue et al. (35) show that the activation of ERK2 by MEK1 is less sensitive to *sevenmaker* mutation: MEK1-facilitated activation activity of mutated *ERK2* is only 0.88 of that for WT *ERK2.* However phosphorylation activity of *ERK2* towards substrate MNK1 is strongly affected by the mutation: phosphorylation activity of mutant *ERK2* is estimated of being 0.11 of that of the *WT* enzyme. This suggests that all the processes involving ERK2 can be negatively affected by *sevenmaker* mutation to a different degree. It also means that the overall balance of these effects *in vivo* is difficult to explicitly estimate since there are many known activators, deactivators and substrates of ERK2 (35), and likely many more will be discovered in the future.

## CONCLUSIONS

Here we developed a kinetic network model to explain the observations of the increased enzymatic activity in the enzymes with the *sevenmaker* mutation and for other similar gain-of-function mutations in the ERK2 enzymes. Our approach presents a comprehensive *quantitative* description of the enzymatic properties of the wild-type and mutated ERK2 regulation systems. First, we constructed a simplified regulation network for ERK2 by arguing that it can be viewed as three coupled Michaelis-Menten processes that describe three main enzymatic processes: activation, deactivation and the phosphorylation. The corresponding kinetic scheme is analyzed then explicitly using the method of first-passage processes to evaluate the enzymatic properties of the system in terms of the individual transition rates and the binding energy between the enzyme and the substrates. The obtained analytical results are also compared with the predictions for the simple Michaelis-Menten scheme. It is argued that mutations modify the interaction energies, and this leads to changes in the enzymatic features of the mutant ERK2 molecules. Our calculations show that the catalytic properties of ERK2 differ significantly from the results for the simplest Michaelis-Menten process. We found a non-monotonic dependence of the specificity constant, which is a quantitative measure of the enzymatic efficiency, as a function of the interaction energy. This suggests that some mutations might increase the activity of the enzyme by changing the interaction energies to the values closer to the observed maximum. The proposed mechanism is also discussed in terms of the fluxes via different branches of the regulation network, and theoretical calculations generally support it. Thus, our main conclusion is that the *sevenmaker* mutation modifies the binding interaction energy in such way that the deactivation process is affected less than the activation processes, leading to the effective increase in the overall catalytic activity. While the mutation lower the rate for each enzymatic branch for some interactions energies, the overall turnover time might at the same time decrease, making them catalytically more active. These theoretical predictions agree with known experimental observations. In addition, it was suggested that the effect of mutation (positive or negative) might depend on the concentration of activator, deactivator and substrate molecules that participate in the ERK2 regulation network.

Our theoretical model provides a consistent chemical description on the possible mechanisms for the gain-of-function mutations in ERK2, giving a fully quantitative measure of mutations, which can be in principle experimentally measured. However, it is important to discuss the limitations of the proposed theoretical method. A weak side of our approach is that a very complex biochemical network with multiple states and transitions, which controls the activities of ERK2 enzymes, is simplified into a network with only three coupled Michaelis-Menten processes. It is also assumed that the reaction constants for activation, deactivation and phosphorylation are the same while they might differ significantly. In addition, current experiments give only very qualitative measurements of the increase enzymatic activities of the mutant ERK2 molecules. But our hope is that the presented quantitative model will stimulate experimental and theoretical studies that will test our ideas, thus advancing our understanding on the mechanisms of functioning of the ERK2 as well as other regulating enzymatic systems.

## AUTHOR CONTRIBUTIONS

A.B.K. designed the research. A.B.K. and M.M. carried out the research, analyzed the data and wrote the paper.

## ACKNOWLEDGMENTS

This work is supported by Center for Theoretical Biological Physics NSF Grant PHY-1427654. A.B.K. also acknowledges the support from Welch Foundation (Grant C-1559) and from the NSF (Grant CHE-1664218).

## LIST OF FIGURES

1 A simplified biochemical regulation scheme for ERK 2 considered in this work. ERK2 first must be phosphorylated by MEK (denoted as *M*) to become an active enzyme and to phoshorylate its substrates (denoted as *S*). At the same time, the phosphatase (denoted as *D*) can dephosphorylate ERK2 to return it to the inactive state. More details are in the text

2 Catalytic constants as functions of the binding energies for Michaelis-Menten (MM, blue) and ERK2 schemes. Negative energies correspond to stronger binding. For MM scheme a monotonic behavior is observed: the stronger the interaction, the lower *k*_*cat*_; while for ERK2 scheme the dependence is non-monotonic: there is an optimal value of binding energy that produces the highest *k*_*cat*_

3 Michaelis constants as functions of the binding energies for Michaelis-Menten scheme (MM, blue) and ERK2 scheme (black). Negative energies correspond to stronger binding. For MM scheme the dependence is monotonic, and the stronger the binding, the lower the Michaelis constant. For ERK2 scheme the dependence is also monotonic, but *K*_m_ changes between two limits

4 Specificity constants (*k*_*cat*_ */K*_m_) as a functions of the binding energy for the Michaelis-Menten scheme (MM, blue) and for the ERK2 scheme (ERK2, black). Negative energies correspond to stronger binding. In the region between two vertical lines for the ERK2 scheme the decrease of strength of interactions will lead to higher values of specificity

5 The specificity constant as the function of the interaction energy for varying contributions from the deactivation process. Numbers in the legend show the concentrations of the deactivator *D* in units of *moles /l.* Dashed line shows the position of the maximum of the specificity constant

6 Inverse catalytic turnover times, or the effective overall reaction rates for the product formation, as a function of the interaction energy for the Michaelis-Menten scheme (MM, blue) and for the ERK2 scheme (black). Concentration of substrate is *S*=10^-5^. In the region between two vertical lines decrease of interaction energies leads to decrease of the reaction rate for MM scheme, but for the ERK2 scheme it leads to the increased turnover rate

